# Towards personalized auditory models: predicting individual sensorineural-hearing-loss profiles from recorded human auditory physiology

**DOI:** 10.1101/2020.11.17.387001

**Authors:** Sarineh Keshishzadeh, Markus Garrett, Sarah Verhulst

## Abstract

Over the past decades, different types of auditory models have been developed to study the functioning of normal and impaired auditory processing. Several models can simulate frequency-dependent sensorineural hearing loss (SNHL), and can in this way be used to develop personalized audio-signal processing for hearing aids. However, to determine individualized SNHL profiles, we rely on indirect and non-invasive markers of cochlear and auditory-nerve (AN) damage. Our progressive knowledge of the functional aspects of different SNHL subtypes stresses the importance of incorporating them into the simulated SNHL profile, but has at the same time complicated the task of accomplishing this on the basis of non-invasive markers. In particular, different auditory evoked potential (AEP) types can show a different sensitivity to outer-hair-cell (OHC), inner-hair-cell (IHC) or AN damage, but it is not clear which AEP-derived metric is best suited to develop personalized auditory models. This study investigates how simulated and recorded AEPs can be used to derive individual AN- or OHC-damage patterns and personalize auditory processing models. First, we individualized the cochlear-model parameters using common methods of frequency-specific OHC-damage quantification, after which we simulated AEPs for different degrees of AN-damage. Using a classification technique, we determined the recorded AEP metric that best predicted the simulated individualized CS profiles. We cross-validated our method using the dataset at hand, but also applied the trained classifier to recorded AEPs from a new cohort to illustrate the generalisability of the method.

## Introduction

Auditory Evoked Potentials (AEPs) are widely adopted as markers of sensorineural hearing loss (SNHL) in clinical and research settings. In research animals, auditory brainstem response (ABR) or envelope-following response (EFR) amplitudes can be used to quantify auditory-nerve (AN) fiber damage, i.e. cochlear synaptopathy (CS), (Kujawa and Liberman, 2009; Furman et al., 2013; Sergeyenko et al., 2013; Shaheen et al., 2015). However, applying the same AEP markers for CS diagnosis in humans has yielded mixed success, since AEP amplitudes can be affected by (i) other coexisting SNHL aspects such as outer-hair-cell (OHC) damage (Don and Eggermont, 1978; Gorga et al., 1985; Herdman and Stapells, 2003; Verhulst et al., 2016; Chen et al., 2008; Garrett and Verhulst, 2019; Keshishzadeh et al., 2020) and (ii) subject-specific factors such as age, gender, and head-size (Trune et al., 1988; Mitchell et al., 1989; Hickox et al., 2017). Moreover, the sensitivity of AEPs to different degrees of OHC-loss and CS is unclear, and a direct quantification of AN fiber damage through histopathology is impossible in live humans (Bharadwaj et al., 2014). These problems hinder the study of the specific impact of OHC-damage and CS on recorded AEPs, and render an AEP-based quantification of AN fiber damage difficult in listeners with mixed hearing pathologies. However, this last step is crucial when developing personalized models of auditory processing for use within numerical closed-loop hearing restoration systems.

Even though several auditory models incorporate sources of SNHL (e.g., Ewert and Dau 2000; Heinz et al. 2001; Rohdenburg et al. 2005; Zilany and Bruce 2006; Jepsen et al. 2008; Jepsen and Dau 2011; Ewert et al. 2013; Verhulst et al. 2018), methods to individualize the AN-damage pattern on the basis of recorded AEP metrics are non-existent. Here, we investigate the potential of common AEP markers to individualize the frequency-specific AN damage profile of personalized auditory models with or without other co-occurring aspects of SNHL. Specifically, we present a combined experimental-modeling method in which (i) individual cochlear-gain-loss (CGL) parameters are extracted from either the audiogram or distortion-product otoacoustic emissions (DPOAEs), and (ii) a feature set of recorded AEP metrics is compared to simulated AEP metrics to derive periphery models with different CS profiles. Using a classifier that was trained on simulated AEPs for different SNHL profiles, we selected the individual AN profile, that best explained the recorded AEP features from a test subject. We tested this method on 35 participants, which were separated into groups of young normal-hearing (yNH), older normal-hearing (oNH) and older hearing-impaired (oHI) listeners (Garrett et al., 2020). Validation of our method to predict individual AN-damage profile from recorded AEPs was performed on data from a new cohort.

Before we describe the classification method in detail, we summarize which AEP markers are promising to include. Among the hitherto proposed AEP-derived metrics of AN damage, the ABR wave-I is known to degrade as a consequence of CS in subjects with intact sensory hair cells (Kujawa and Liberman, 2009; Parthasarathy and Kujawa, 2018), however this metric is highly variable in humans (Stamper and Johnson, 2015; Plack et al., 2016) when the contribution of between-subject variability sources such as head-size or tissue resistance are not considered (Prendergast et al., 2018). Even though we can assume that any hearing deficit reflecting on the ABR wave-I would travel through the auditory pathway to reflect on the ABR wave-V as well, homeostatic gain changes between AN fibers and inferior colliculus (IC) may affect the wave-V amplitude (Schaette and McAlpine, 2011; Chambers et al., 2016; Möhrle et al., 2016; Henry and Abrams, 2018) and hence its diagnostic power for CS diagnosis. Another AEP marker, the EFR amplitude, which reflects the strength of a phase-locked AEP response to an amplitude-modulated (AM) stimulus, was shown to degrade as a consequence of CS in mice histological studies (Shaheen et al., 2015; Parthasarathy and Kujawa, 2018) and as a consequence of age in human listeners (Goossens et al., 2016; Vasilkov et al., 2020). EFRs offer a more robust measure of the AN fiber population than the ABR wave-I, when recorded in the same animals (Shaheen et al., 2015; Plack et al., 2016; Parthasarathy and Kujawa, 2018). However, similar to the ABR wave-V, EFR generators have latencies associated with IC processing (Purcell et al., 2004), thus differences in central auditory processing may reflect on the EFR magnitude to mask individual synaptopathy differences (Chambers et al., 2016; Möhrle et al., 2016; Parthasarathy et al., 2019a,b). To address these issues, relative EFR and ABR metrics were proposed in several studies to cancel out subject-specific factors and isolate the CS component of SNHL in listeners with coexisting OHC-loss: ABR wave-I amplitude growth as a function of stimulus intensity (Furman et al., 2013), ABR wave-I -V latency difference (Coats and Martin, 1977; Elberling and Parbo, 1987; Watson, 1996), the wave-V and I amplitude ratio (Gu et al., 2012; Schaette and McAlpine, 2011; Hickox and Liberman, 2014), EFR amplitude slope as a function of modulation depth (Bharadwaj and Shinn-Cunningham, 2014; Guest et al., 2018), the derived-band EFR (Keshishzadeh et al., 2020), or the combined use of the ABR wave-V and EFR (Vasilkov and Verhulst, 2019). While these relative metrics are promising, it is not known how OHC-loss and CS differentially impact AEPs. Recent modelling approaches have shown promise to design EFR stimuli which are maximally sensitive to CS in the presence of OHC damage (Vasilkov et al., 2020), but conclusive histopathological evidence is to date not available. To make use of the listed metrics to build personalized hearing profiles for a broad population with various SNHL etiologies, two requirements need to be fulfilled. We need to (i) use AEP markers that are maximally sensitive to the CS aspect of SNHL and (ii) combine them with a sensitive marker of OHC deficits to individualize the OHC and CS aspects of SNHL. We thus considered various AEP markers (a total of 13) encompassing spectral magnitudes, time-domain peaks, latencies and relative metrics, and combinations thereof, to identify which markers best predict the simulated individualized CS profiles and can be used for reliable auditory profiling.

### Experimental Design

ABR, EFR and OHC-damage markers were derived from recordings of two experimental setups in different locations. These recordings were used for development and validation of the proposed method, respectively.

#### Participants

The dataset that was used to develop the auditory profiling method included recordings from a total of 43 subjects. They were recruited into three groups: 15 young normal-hearing (yNH: 24.53±2.26 years, 8 female), 16 older normal-hearing (oNH: 64.25±1.88 years, 8 female) and 12 older hearing-impaired (oHI: 65.33±1.87 years, 7 female) groups. Two oNH subjects were omitted from our study due to non-identifiable ABR waveforms. The hearing thresholds of the participants were assessed at 12 standard audiometric frequencies between 0.125 and 10 kHz (Auritec AT900, Hamburg, Germany audiometer). AEP stimuli were presented monaurally to the ear with the best 4 kHz threshold. Audiometric thresholds were below 20 dB-HL at all measured frequencies in the yNH group and below 25 dB-HL for frequencies up to 4 kHz in the oNH group. The oHI listeners had sloping high-frequency audiograms with 4-kHz thresholds above 25 dB-HL (Fig. 1a). The AEP recordings were conducted in an electrically and acoustically shielded booth, while subjects were sitting in a comfortable chair and watching silent movies.

**Figure 1:**
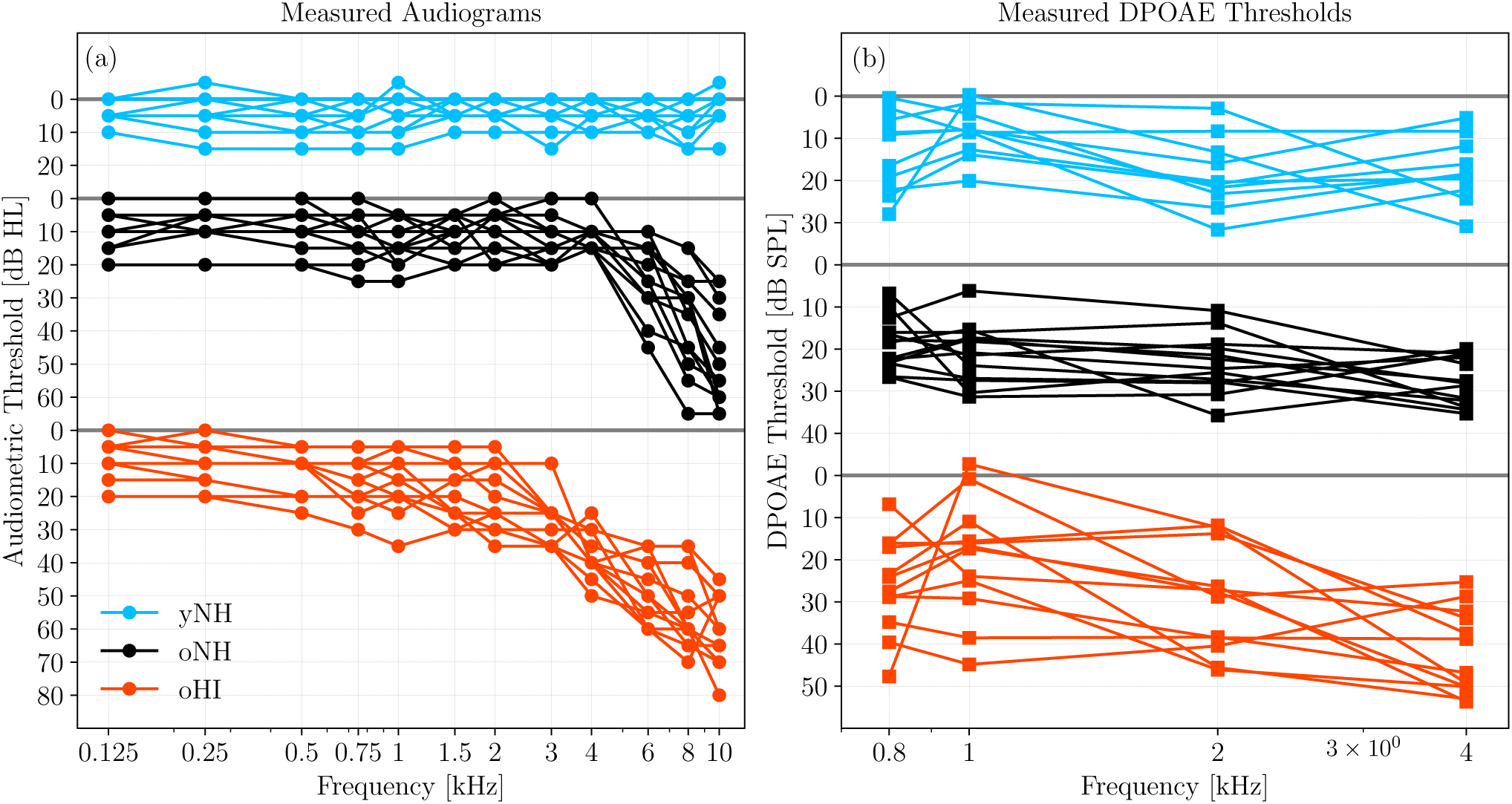
(a) Audiograms and (b) DPOAE thresholds (DPTHs) of the participants in the first experiment.

The second experiment, which was used to validate our method on a new cohort, had 19 yNH subjects, aged between 18 and 25 years (21.6±2.27 years, 12 female). Volunteers with a history of hearing pathology or ear surgery were excluded based on a recruitment questionnaire. Audiograms were measured in a double-wall sound-attenuating booth, using an Interacoustics Equinox Interacoustics audiometer. All participants had audiometric thresholds below 25 dB-HL within the measured frequency range, i.e. [0.125-10] kHz, and the best ear was determined on the basis of their audiogram and tympanogram. The experiment protocol included AEP measurements with a maximum duration of 3 hours and we only considered one AEP metric for validation purposes in the present study. AEP recordings were conducted in a quiet room while subjects were seated in a comfortable chair and watching muted movies. To minimize the noise intrusion level, both ears were covered with earmuffs and all electrical devices other than the measurement equipment (Intelligent Hearing Systems) were turned off and unplugged.

Participants of both experiments were informed about the experimental procedure according to the ethical guidelines at Oldenburg University (first experiment) or Ghent University Hospital (UZ-Gent, second experiment) and were paid for their participation. A written informed consent was obtained from all participants.

#### Distortion Product Otoacoustic Emission (DPOAE)

In the first experiment, DPOAEs were acquired and analyzed using a custom-made MATLAB software (Mauermann, 2013). Stimuli were delivered through ER-2 earphones coupled to the ER-10B+ microphone system (Etymotic Research) using a primary frequency sweeping procedure at a fixed f_2_*/*f_1_ ratio of 1.2. The implemented DPOAE paradigm, continuously swept the primary frequencies with a rate of 2s/octave within a 1/3 octave range around the geometric mean of f_2_ ∈ {0.8, 1, 2, 4} kHz (Long et al., 2008). The L_2_ primary levels ranged between 30-60 dB-SPL for the yNH and oNH groups, using a 6-dB step. The level range was different for the oHI group: 30-72 dB-SPL. L_1_ levels were determined according to the scissors paradigm (Kummer et al., 1998). For a given f_2_ primary frequency, the DP-component (L_DP_) growth function was plotted as a function of L_2_ and a cubic curve was fit to the L_DP_ data-points using a bootstrapping procedure to include the standard deviation of the individual L_DP_ data-points in the fit (Verhulst et al., 2016). The L_2_ level at which the cubic curve crossed −25 dB-SPL was determined for each bootstrap average to yield the DPOAE threshold (DPTH) and its standard deviation at a given f_2_ (Boege and Janssen, 2002). Derived experimental DPTHs of the yNH, oNH and oHI groups are shown in Fig 1b. DPOAEs were not available for the subjects of the validation experiment.

#### EEG Measurements

ABR and EFR stimuli were generated in MATLAB and digitized with a sampling rate of 48 kHz for the first dataset. Afterwards, they were delivered monaurally through a Fireface UCX external sound card (RME) and a TDT-HB7 headphone driver connected to a shielded ER-2 earphone. The electroencephalogram (EEG) signals were recorded with a sampling frequency of 16384 Hz via a 64-channel Biosemi EEG system using an equidistantly-spaced electrode cap. All active electrodes were placed in the cap using highly conductive gel. The common-mode-sense (CMS) and driven-right-leg (DRL) electrodes were attached to the fronto-central midline and the tip of the nose, respectively. A comprehensive explanation of the experimental configuration can be found in Garrett and Verhulst (2019).

AEPs of the validation experiment were recorded using the SmartEP continuous acquisition module (SEPCAM) of the Universal Smart Box (Intelligent Hearing System, Miami, FL, United States). EFR stimuli were generated in MATLAB using a sampling rate of 20-kHz and stored in a “.*wav*” format. These files were loaded in SEPCAM and converted to the “.*stm*”, SEPCAM compatible format. AEP stimuli were presented monaurally through a shielded ER-2 earphone (Etymotic Research) and AEPs were recorded at a sampling frequency of 10 kHz via Ambu Neuroline 720 snap electrodes connected to vertex, nasion and both earlobes. The electrodes were placed after a skin preparation procedure using NuPrep gel. The skin-electrode impedance was kept below 3 kΩ during the recordings.

#### EFR stimuli

We recorded EFRs in response to a 400-ms-long stimuli consisting of a 4-kHz pure-tone carrier and a 120-Hz rectangular-wave modulator with 25% duty cycle (i.e. the RAM25 in Vasilkov et al. 2020). The stimulus waveform is visualized in the inset of Fig. 2b and we considered a modulation depth of 95%. Stimuli were presented 1000 times (500 times in either positive or negative polarity) and had a root-mean-square (RMS) of 68.18 dB-SPL. The calibration of the stimulus was performed to have the same peak-to-peak amplitude as a 70-dB-SPL sinusoidal amplitude modulated (SAM) 4-kHz pure-tone. The Cz channel recording was re-referenced to the average of the ear-lobe electrodes and 400-ms epochs were extracted relative to the stimulus onset. The mean-amplitude of each epoch was subtracted to correct for the baseline-drift. See Vasilkov et al. (2020) for further details on the frequency-domain bootstrapping and noise-floor estimation method. The noise-floor corrected spectral magnitudes 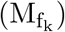 at the modulation frequency f_1_ = 120Hz and four harmonics, i.e., f_2_ to f_5_, were summed up to yield the EFR.

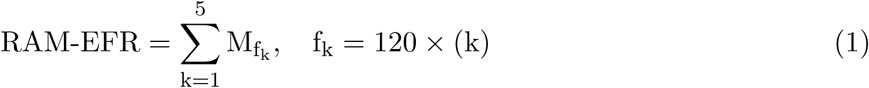

**Figure 2:**
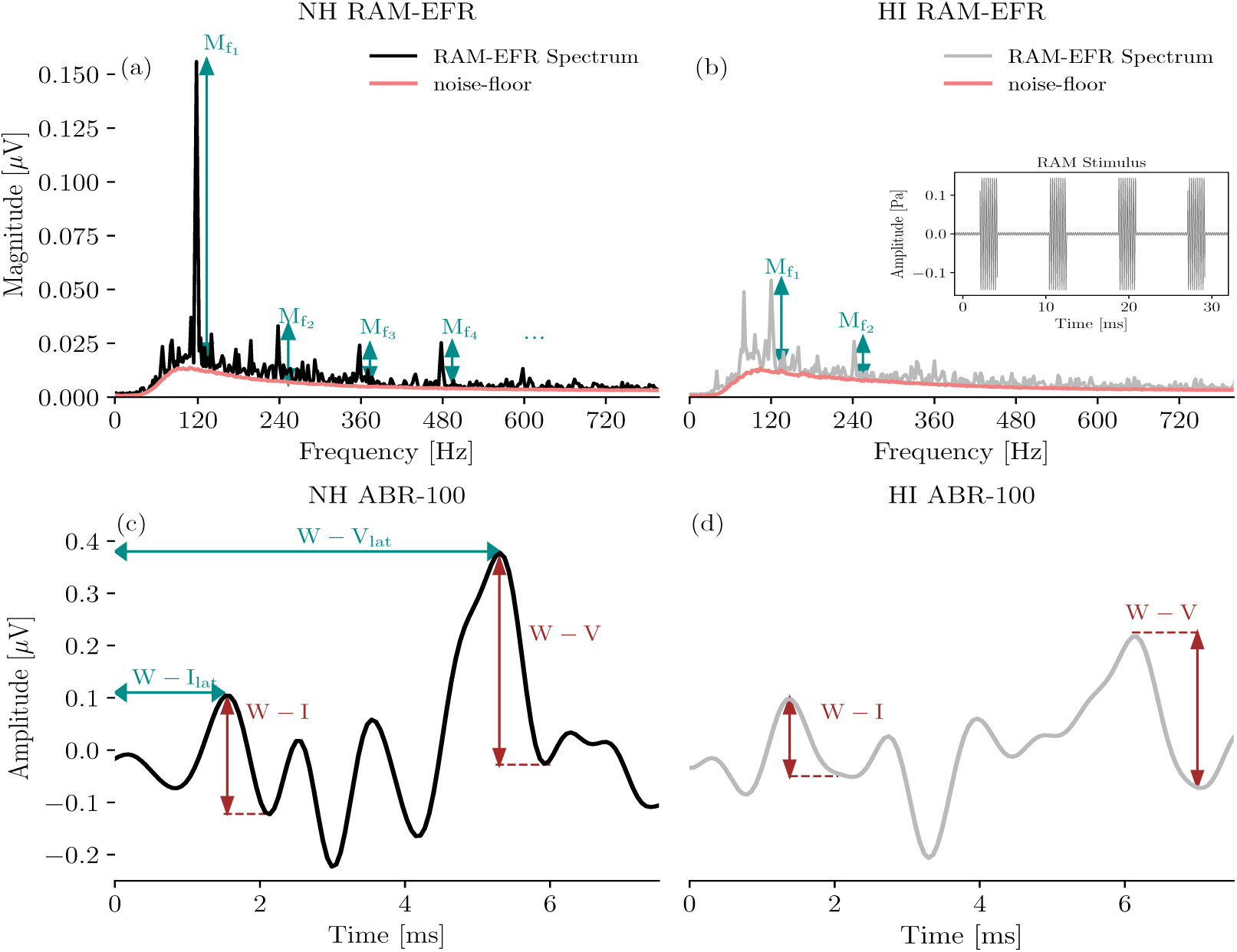
Comparison of exemplary NH and HI RAM-EFRs and ABRs. (a) RAM-EFR of a yNH subject (yNH15) and the corresponding noise-floor (NF). Arrows specified by M_f_, show the peak-to-noisefloor magnitudes at the modulation frequency, i.e., 120 Hz, and the following harmonics. (b) RAM-EFR of an oHI subject (oHI) and the corresponding NF. (c) ABR of a yNH subject (yNH15). Arrows show the extracted wave-I and V amplitudes and latencies. (d) ABR of an oHI subject (oHI12).

Figure 2a depicts a typical NH RAM-EFR spectrum and corresponding noise-floor. The arrows show the derived peak-to-noise-floor magnitudes at the modulation frequency and following harmonics. The energy of EFR peak is reduced for the oHI subject shown in the panel (b).

The RAM-stimulus in the second experiment (i.e. the validation database) was a 110-Hz 95% modulated 4-kHz pure-tone. The 500-ms stimulus was presented 1000 times with alternating polarity (500 each) and had a 70 dB-SPL level. The acquired AEPs were initially saved in “.*EEG*.*F*” format on SEPCAM and were afterwards converted to “.mat” format using the custom-made “*sepcam2mat*” MATLAB function for offline analysis. EFRs recorded from the vertex electrode were re-referenced to the ipsilateral earlobe electrode and filtered between 30 and 1500 Hz using an 800^th^ order Blackman-window based finite-impulse-response (FIR) filter. Epoching was applied to the steady state part of the response, i.e. 100 to 500 ms of the response relative to the stimulus onset. The baseline drift was corrected by subtracting the mean of each epoch, afterwards 200 epochs with the highest peak-to-trough values were rejected. The amplitudes of the remained epochs did not exceed 100 *µ*V. A frequency-domain bootstrapping approach was adopted to estimate the noise-floor and to remove it from the averaged trials using the method proposed in Zhu et al. (2013). To this end, we calculated the fast Fourier Transform (FFT) of 800 epochs to generate 400 mean spectra by randomly sampling the 800 epochs with replacement (keeping an equal number of polarities in the draw). Averaging the resampled spectra yielded the *i-th* mean-EFR spectrum 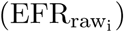:

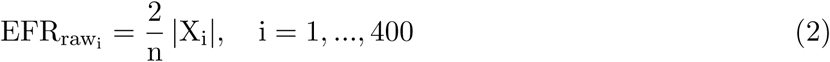

where, X_i_ stands for the *i-th* averaged resampled spectra and n is the number of FFT points (n=10000). To calculate the spectral noise-floor, we repeated the resampling procedure 1500 times, but used phase-flipped odd epochs:

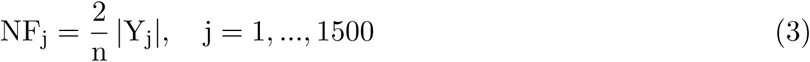

In Eq. 3, Y_j_ is the *j-th* averaged resampled spectra with phase-flipped odd epochs. Lastly, we subtracted the NF mean 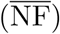, from each of the 400 bootstrapped mean-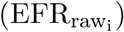 to derive 400 NF-corrected EFR spectra:

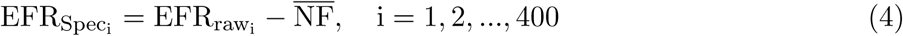

The peaks of the 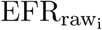 at the modulation frequency of stimulus (f_1_=110 Hz), and the next four harmonics were identified if they were above the 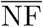. We defined the RAM-EFR_i_ by summing the magnitudes of the identified peaks for each 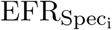 The RAM-EFR metric mean and variability was defined by the mean and standard deviation of RAM-EFR_i_ over 400 samples.

#### Auditory Brainstem Responses

ABRs were recorded to 80-*µ*s-long alternating polarity clicks presented at 70 and 100 dB-peSPL. Stimuli were presented through the setup explained in Garrett et al. (2019) and repeated 3000 times with a rate of 10 Hz using a uniformly distributed random inter-stimulus interval of 100 ms±10 ms. Cz-channel recordings were re-referenced to the contra-lateral earlobe electrode and filtered between [100-1500] Hz. 25 ms-long epochs, i.e. −5 to 20-ms relative to the stimulus onset, were extracted and corresponding mean values were subtracted to perform a baseline correction. Then, each positive polarity epoch was averaged with the following negative epoch and 100 paired-averages with the highest peak-to-trough values were rejected. The remaining pair-averaged epochs had amplitudes below 25 *µ*V. To include ABR variability in our analysis and to estimate the ABR noise-floor, we applied the bootstrapping approach of Zhu et al. (2013), in the time domain. 2000 and 4500 epochs were drawn for the signal and noise-floor estimation, respectively. Half of the noise-floor-estimation epochs (i.e. 2250 pair-averaged drawn epochs with replacement) were multiplied by −1 before final averaging. Finally, the estimated noise-floor mean was subtracted from the 2000 averaged epochs to yield mean noie-floor-corrected ABR waveforms. ABR wave-I and -V peak and trough amplitudes and corresponding latencies were determined by visual inspection from the mean ABR waveform and were confirmed by an audiologist. Figure 2 (panels c and d) compares ABR waveforms of a yNH and oHI subject from the cohort and indicates the identified ABR peaks and latencies. To extract peak latencies and amplitudes from the bootstrapped data, wave maxima and minima were detected in 1, 1.8, 0.5 and 1.5 ms intervals around the wave-I_70_, wave-V_70_, wave-I_100_, wave-V_100_ peaks and troughs identified from the mean ABR waveform. The interval ranges were determined based on visual inspection. ABR wave-I and V latencies were shifted by 1.16 ms to compensate for the delay introduced by the sound-delivery system.

We used a total of 13 ABR and EFR markers in the development phase and one EFR marker in the validation phase. Table 1 details the definition of each metric and lists the corresponding abbreviations used in this paper. The last column defines the variability metric associated with each marker, which were obtained from the earlier described bootstrapping procedure. To determine the measurement variability of ABR growth-slopes, we applied error propagation to account for the standard deviations of two different measures from the same listener, e.g., ABR-70 and ABR-100. In this case, the bootstrapped metrics were drawn from the 95% confidence interval of a normal distribution characterized by the mean of the metric and its bootstrapped standard deviation. The bootstrapping technique described in this section, provided a tool to estimate the variability of AEP-derived metrics and to incorporate them in the proposed classification approach. Obtained standard deviations from bootstrapping can be used to measure the CS-profiling prediction robustness of the study participants.

**Table 1:**
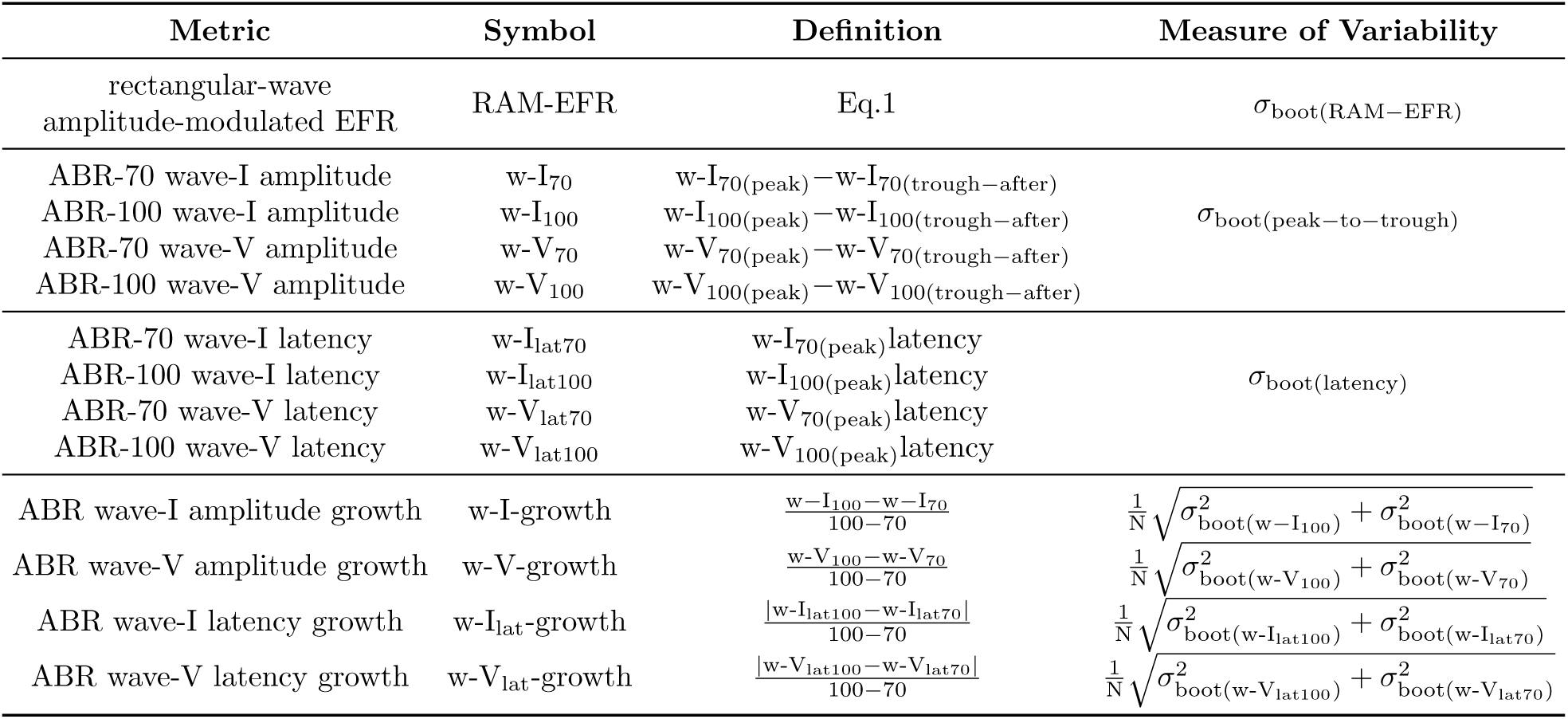
Extracted AEP-metrics definitions and corresponding standard deviations. In the last column, *σ* represents the standard deviation. *σ*_boot_ is the standard deviation of the bootstrapped metric.

### Individualized Auditory Periphery Model

To simulate individualized SNHL profiles that would match the histopathology of the study participants, we used a computational model of the auditory periphery (Verhulst et al., 2018; Osses and Verhulst, 2019). In the first step, we personalized the cochlear model parameters on the basis of OHC markers of SNHL (audiogram or DPTH). Afterwards, we simulated AEPs for different degrees of CS and compared the simulations to the recordings to develop and test our auditory profiling method. Figure 3 schematizes the auditory model individualization.

**Figure 3:**
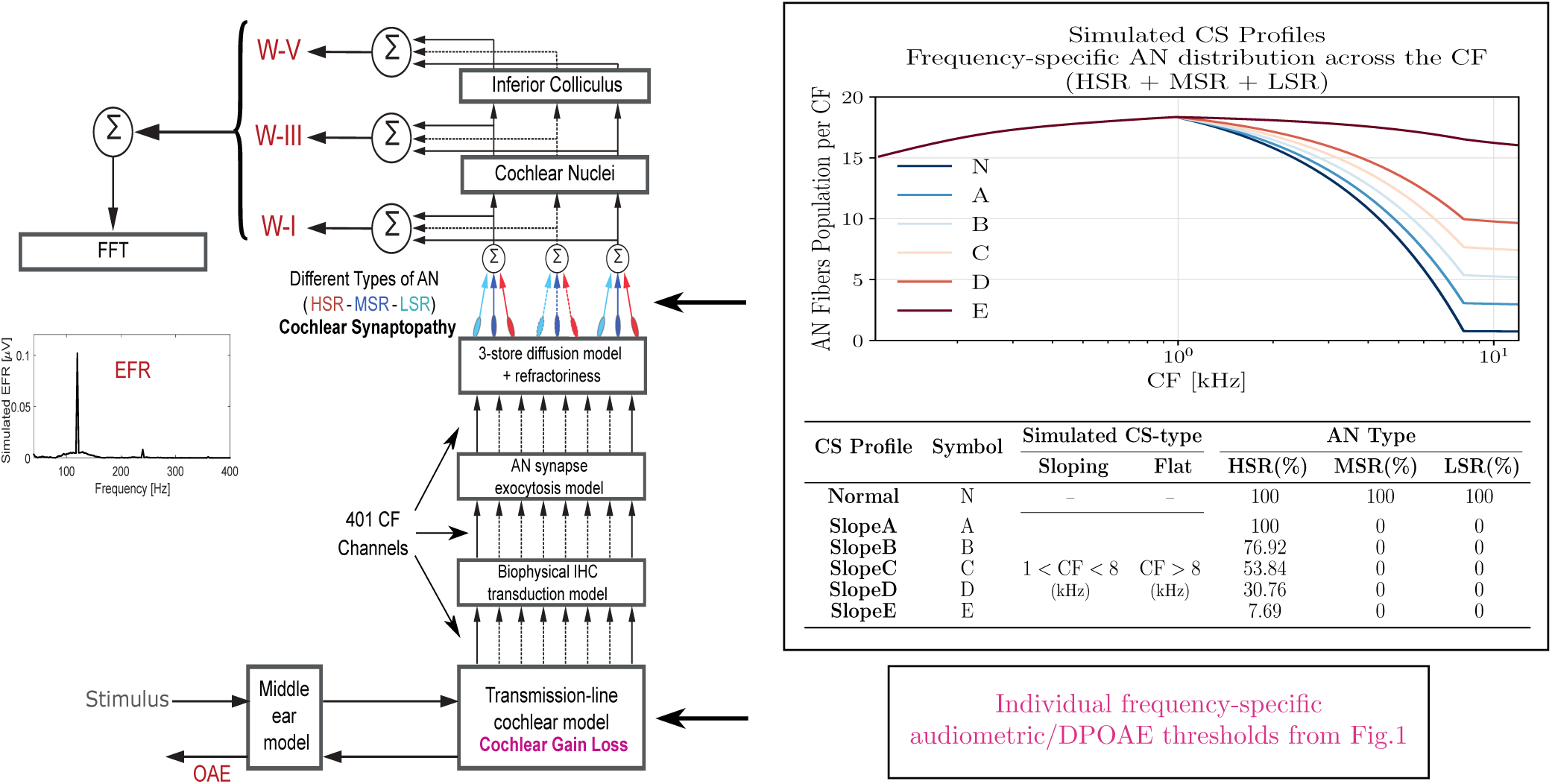
Auditory model individualization. The block-diagram on the left depicts the different stages of the employed auditory periphery model (Verhulst et al., 2018). Experimentally measured audiometric thresholds were inserted to the transmission-line cochlear model to adjust BM admittance function poles. The box on top-right corner, shows the non-uniform AN population distribution across the CF for simulated different degrees of CS profiles. Ti he profile without CS is shown in dark brown (indicated with N) and higher degrees of CS are shown according to the color-map.

#### Cochlear Model Individualization

Measured audiograms and DPTHs were used independently to determine the individual CGL parameters (in dB-HL) of the cochlear transmission-line (TL) model, shown in pink in Fig.3. In our approach, CGL determines the double-pole of the cochlear admittance through the gain and tuning of the cochlear filters (Verhulst et al., 2012). We thus model the consequence of OHC-damage or presbycusis without specifically accounting for damage of the stereocilia or sensory cells. From here on, mAudTH and sAudTH refer to measured and simulated audiometric thresholds, respectively. Likewise, mDPTH and sDPTH stand for measured and simulated DPOAE thresholds.

#### Audiogram-based cochlear filter pole-setting

Here, we translated the frequency-specific audiometric dB-HL (Fig. 1a) into cochlear filter gain loss. These values were translated into double-pole values of the cochlear admittance function across CF (see Verhulst et al. 2016).

Specifically, at a CF corresponding to a measured audiometric frequency (CF = f_aud_), the power spectrum of the NH basilar membrane (BM) impulse response, H_NH_(f_aud_), served as reference before the gain loss was applied. Among a range of cochlear filter pole-values in [0.036,0.302], the polevalue, 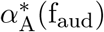, that causes a relative gain-loss equal to mAudTH(f_aud_), was assigned. Thereby, the CGL at CF = f_aud_ is given by:

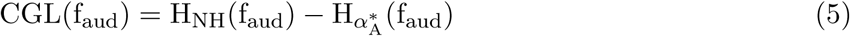

where 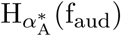 equals the power spectrum of the BM impulse response at CF = f_aud_ with a pole value of 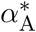 that causes a CGL equal to mAudTH(f_aud_). This procedure was repeated for all CF channels corresponding to measured audiometric frequencies and individualized cochlear filter pole-functions were obtained by interpolating the pole-values across CF (Verhulst et al., 2016).

We employed the predicted pole functions to simulate individual audiograms and to evaluate the prediction error. To this end, individualized AN excitation patterns (ANEP) were simulated in response to 500-ms pure-tones presented at audiogram frequencies (f_aud_) using 62 intensity levels (L) between −5 and 55 dB-SPL. We defined ANEP as the RMS of the AN firing rate at each CF ∈ f_aud_ and determined on-CF peaks of the presented level series as 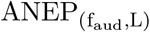. We simulated NH ANEPs using NH pole-function at the threshold of audibility in a frequency-specific manner (L_NH_(f_aud_)), i.e. the zero-phon curve of the equal-loudness-contour (ISO 226:1987). From this reference NH curve, we calculated the simulated audiometric thresholds (sAudTH) of the experiment participants as follows:

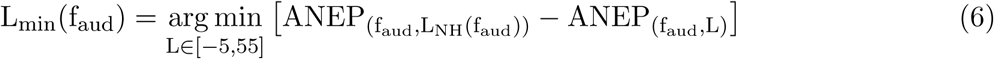

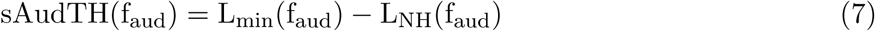

Figure 4a shows grand-averaged mAudTHs and sAUDTHs across the yNH, oNH and oHI groups. Additionally, Fig. 4c, compares the sAudTH (dashed lines) and mAudTH (solid lines) of an example yNH and oHI subject. Note that simulating CGLs greater than 35 dB-HL is impossible in our cochlear model, which has a maximal applicable cochlear mechanical gain of 35 dB. In the last step, we estimated the absolute prediction error as follows:

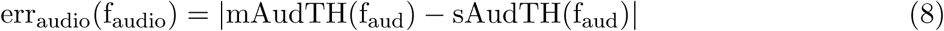

**Figure 4:**
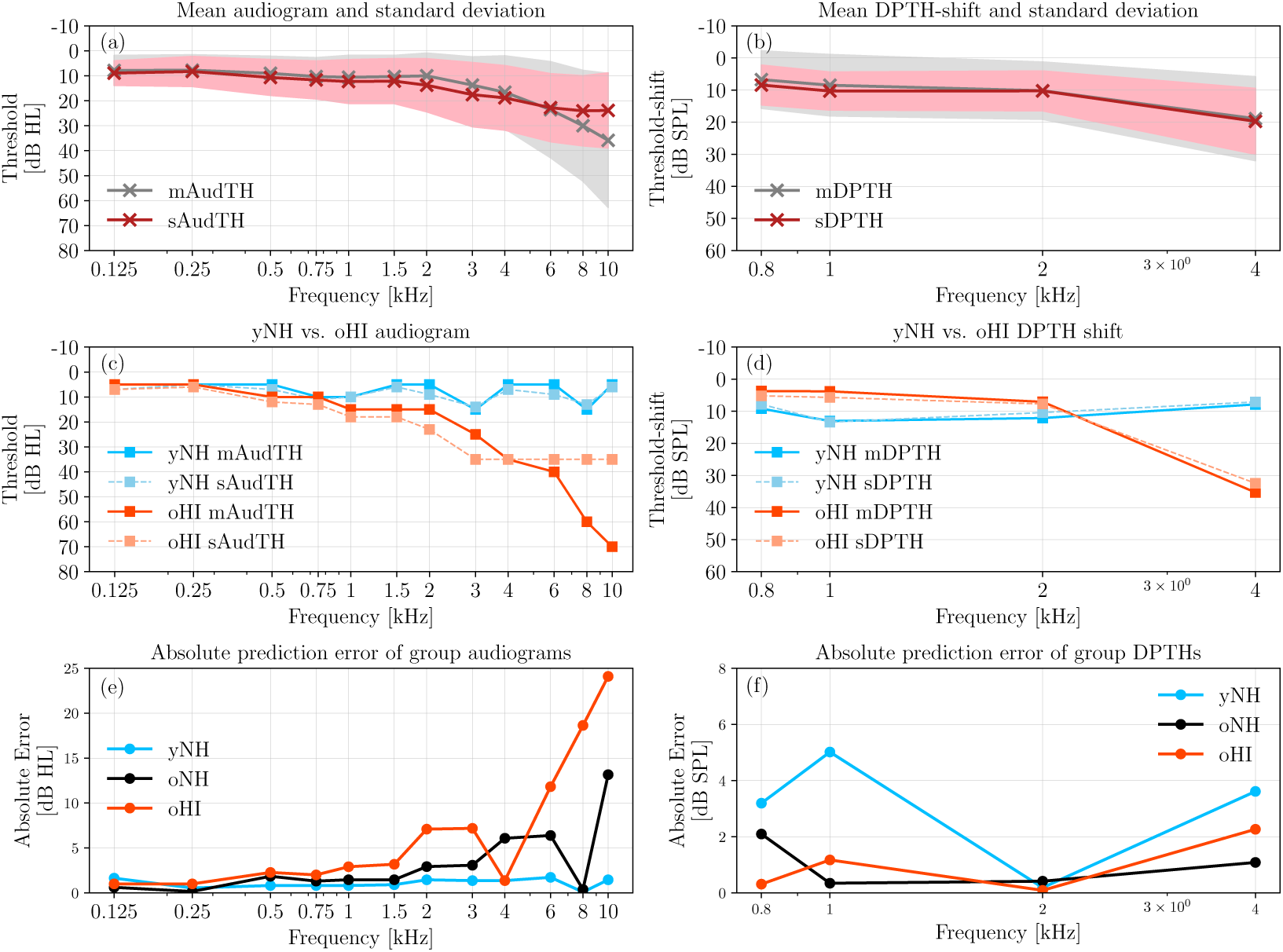
A comparison between the measured and simulated AudTHs and DPTHs. The average (solid) and standard deviation (shaded area) of the measured (grey) and simulated (red) AudTHs and DPTHs are shown in panel (a) and (b), respectively. A comparison between sAudTH and mAudTH of a yNH and oHI listener is shown in panel (c). Panel (d) compares the sDPTH (dotted) and mDPTH (solid) of the same yNH and oHI listeners (c). Frequency-specific group-averaged absolute prediction errors of AudTH and DPTH are shown in panels (e) and (f), respectively (yNH: blue, oNH: black, oHI: orange).

Figure 4e compares the mean absolute errors on a group-level basis. The elevated error of the oHI group at high frequencies is due to the model limitation in simulating gain losses greater than 35 dB-HL.

#### DPTH-based cochlear filter pole-setting

Implementing DPTH-based cochlear model individualization was complicated by the fewer DPTHs we had available, i.e. four frequencies, compared to 12 AudTHs. Hence, a simple interpolation to determine poles between the measured frequencies, yielded large prediction errors. Additionally, the longitudinal filter coupling and associated gain propagation along the cochlear partition complicated matters. To tackle these issues, we trained a machine-learning algorithm to map DPTHs via cochlear travelling waves to corresponding cochlear filter pole functions across CF. Once trained, we need only a few measured DPTHs to make a relatively accurate prediction of individual pole values. Figure 5 illustrates the complete procedure.

**Figure 5:**
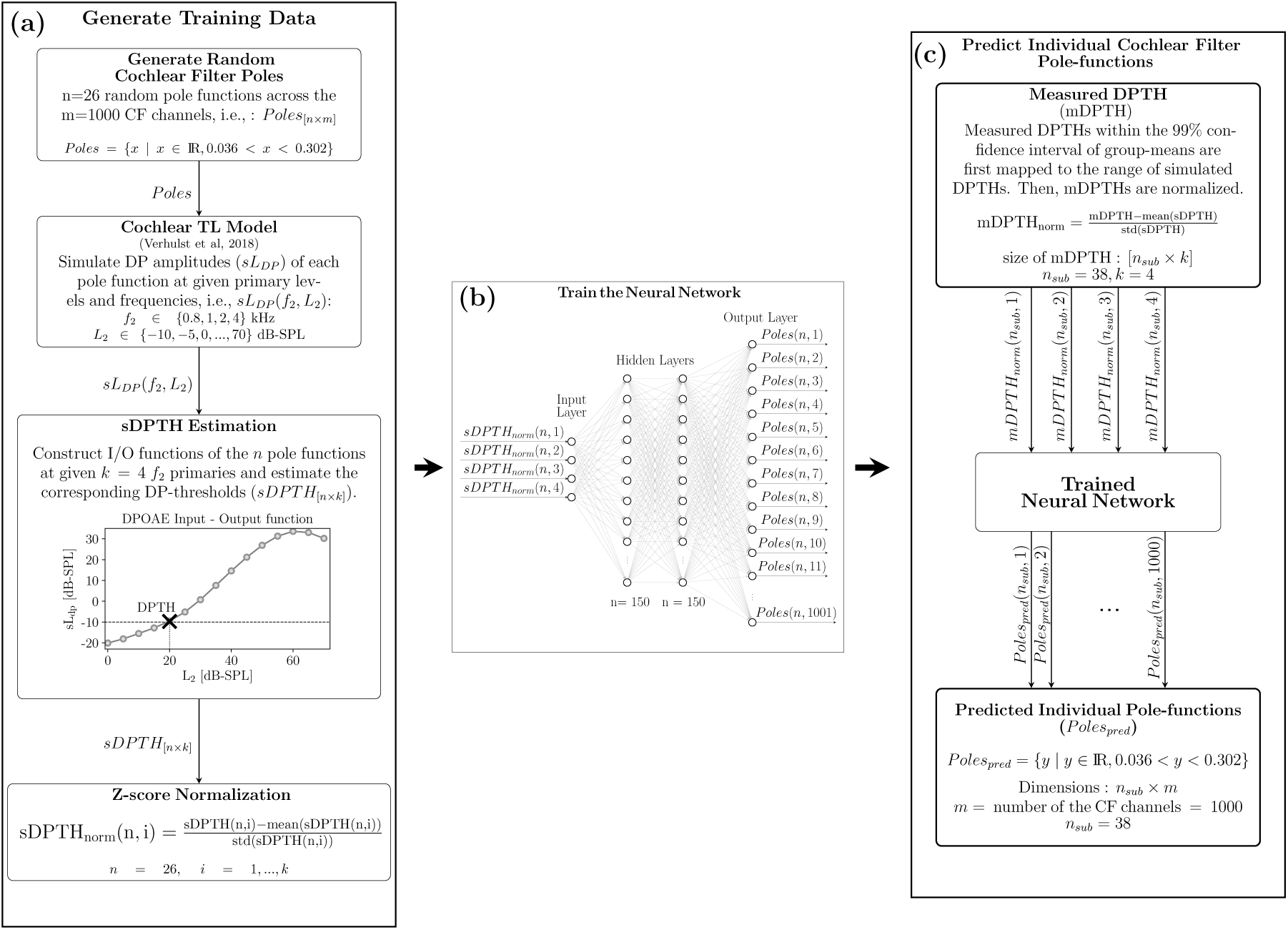
Neural network-based cochlear model individualization using measured and simulated DPTHs. (a) Random cochlear filter poles are generated and corresponding DPTHs are simulated using TL model (sDPTH). (b) The normalized sDPTH (sDPTH_norm_)at four frequencies are introduced to the neural-network as input. The random pole values generated in (a) are served as training target for sDPTH_norm_. (c) Measured DPTHs (mDPTHs) are fed into the trained neural network after pre-processing and individualized cochlear filter pole-functions are predicted.

First, we constructed the training data (Fig. 5a) using 26 sets of random cochlear filter pole functions. Each set contained 1001 CF-dependent poles and random pole-values lay between 0.036 and 0.302, covering the pole-values associated with both NH and HI profiles. Additionally, three reference pole-functions were included as part of the training: NH_poles_ (NH poles), flat_min_ with across-CF poles of 0.036 (maximally intact cochlea) and flat_max_, with across-CF poles of 0.302 (35 dB-HL across CF). We employed the generated pole functions and simulated DP amplitudes (sL_DP_: the magnitude of 2f_1_ *−* f_2_) to train the mapping function. The considered f_2_ primary frequencies, i.e. 0.8, 1, 2 and 4 kHz (f_1_ = f_2_*/*1.2) corresponded to the recordings we had available and L_2_ levels (−10 to 70 dB-SPL, with a step of 5-dB). We simulated DPOAE input-output (I/O) functions at each f_2_ frequency and determined the sDPTH as the L_2_ level at which the sL_DP_ growth function crossed the L_2_ of −10 dB-SPL. We chose a −10 dB-SPL threshold for our simulations, given that the conventional experimental −25 dB-SPL crossing point yielded inconclusive sDPTH, in particular for pole values associated with greater CGLs. sDPTH values for 26 sets of pole-functions at four primary frequencies were fed into the neural network after normalization (sDPTH_norm_, Fig. 5b) to train it to map frequency-specific sDPTH_norm_ values (input) to CF-dependent pole-functions (output).

The architecture of the designed neural network is shown in Fig. 5b, and consists of an input-layer of four neurons, two hidden-layers of 150 neurons and an output layer of 1001 neurons. A standard *sigmoid* activation function (i.e., between 0 and 1), was applied to the hidden layers. A customized *sigmoid* activation function (between 0.036 to 0.302), was employed in the output layer to yield the desired range of the cochlear model pole-functions. An ADAM optimizer with a learning rate of 0.001 was applied to minimize the mean-square-error (MSE) of the learning algorithm. The method was developed in Python using Keras library and Tensorflow back-end.

The trained neural network was employed to predict individualized pole-functions given DPTHs of the experimental cohort (Fig. 5c). Prior to the prediction, mDPTHs needed to be pre-processed to determine a suitable experimental range of DPTHs for the mapping. Among the 41 subjects, six subjects (yNH: three, oNH: two and oHI: one) without complete mDPTH values at all measured frequencies were dropped. In each of the three recruited groups, the 99% confidence interval around the frequency-specific group-means were specified and mDPTH values that either exceeded or fell below of those intervals were set to extremum values. Then, mDPTHs were mapped to the range of the sDPTH associated with reference flat_min_ 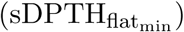 and flat_max_ 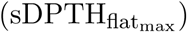 pole functions. Afterwards, mapped mDPTHs (mDPTH_map_) were normalized (mDPTH_norm_) and given to the trained neural network to predict personalized pole-functions. To assess the prediction error, the predicted pole functions (Poles_pred_ in Fig. 5c), were used to simulate individualized sDPTHs that were compared to the individual mDPTHs f_2_ primary frequencies. mDPTHs and sDPTHs were referenced to the simulated DPTHs of a model with NH_poles_ as follows:

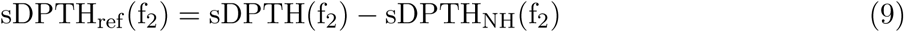

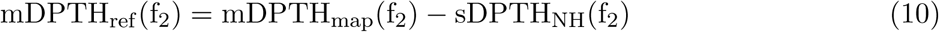

sDPTH_NH_(f_2_) refers to the frequency-specific sDPTH values simulated using the model with NH_poles_. Obtained sDPTH_ref_ and mDPTH_ref_ from Eq. 9 and 10 were mapped back to the experimental range according to Eq. 11 and 12, and corresponding grand-averages and standard deviations are shown in Fig. 4b. More specifically, Fig. 4c compares measured and simulated DPTH-shifts for a yNH and oHI subject.

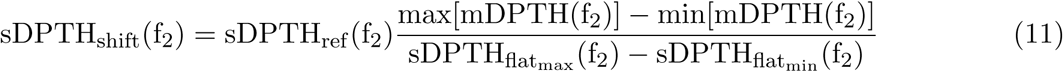

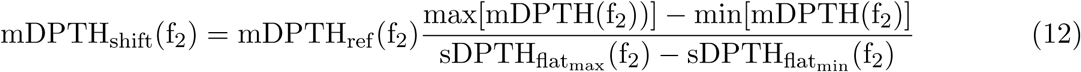

Lastly, the prediction error was calculated as in Eq. 13 and the absolute mean error for each group is shown in Fig 4f.

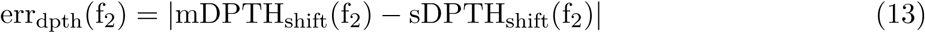

The developed machine-learning approach can be used to personalize cochlear model parameters based on an objective measure of OHC damage (DPTH) and predict individual CS profiles. CS-profiling can be compared for either the DPTH or AudTH-based cochlear model individualization method, and when no DPTHs are available the standard audiogram-based method can be adopted.

### Simulating Cochlear Synaptopathy Profiles

We employed the AudTH- and DPTH-based individualized CGL models to simulate EFRs and ABRs for different CS profiles. To introduce CS, the simulated normal-hearing AN fiber populations, (the *N* CS profile in Fig. 3) was reduced in a CF-specific manner. Five additional CS profiles were simulated by proportionally lowering the number of different AN types, starting from low- and medium-spontaneous-rate (LSR and MSR) fibers in profile *A* to the most severe AN-loss in *E* that only kept 7.69% of the high-spontaneous-rate (HSR) fiber population. The table in Fig. 3 details the AN fiber numbers and types considered for each of the six simulated CS profiles. IHC-related dysfunctions were not considered in this study, given that low degrees of CS do not cause IHC damage (Kujawa and Liberman, 2009; Furman et al., 2013; Shaheen et al., 2015).

However, removing all AN fibers from an IHC in the model would functionally correspond to IHC damage. The CF dependence of the AN population was considered in two steps: (1) Following the CF-dependent AN distribution observed in rhesus monkey (Valero et al., 2017; Keshishzadeh et al., 2020), we applied a non-uniform NH AN fiber population. (2) CF-specific AN-damage profiles were simulated. The former was achieved by mapping the counted CF-dependent AN fibers population in the rhesus monkey (Valero et al., 2017) to the human cochlea, using a distribution of N_HSR_ = 68%, N_MSR_ = 16% and N_LSR_ = 16% at each CF (Liberman, 1978). Then, sloping high-frequency AN-fiber loss was applied across CF with the assumption that CS starts from the higher frequencies first (Wu et al., 2020). We ran EFR/ABR simulations for different AN fiber damage profiles, which were characterized by a sloping loss of between 1 and 8-kHz. Above 8 kHz, we applied a frequency-independent loss.

For every subject we simulated AEPs for each CS profile, after we personalized the cochlear models using either the AudTH-or DPTH-based method. The stimuli adopted for these simulations were identical to those adopted experimentally, but were digitized using a sampling rate of 100 kHz, rather than 48 kHz. Simulated instantaneous firing rates from the AN, cochlear nucleus (CN) and IC model stages, namely ABR wave-I, III and V, respectively, were added up to simulate EFRs (Fig.3). RAM-EFR magnitudes were calculated using Eq. 1.

To simulate ABRs, 80-*µ*s clicks were presented to the model with a continuous sequence of 50 repetitions of alternating polarities (100 in total) and a rate of 10 Hz. Sequential stimulus presentation was adopted to account for the adaptation properties of AN fibers. Individual ABR wave-I and V latencies and amplitudes were extracted by averaging the peak-to-trough values of the response to the last, i.e. 50^th^, positive and negative clicks. The simulated ABR wave-I and V latencies were respectively shifted by one and three ms to match latencies of recorded ABRs. These values were determined to match the measured yNH group-mean ABR wave-I and V latencies (at 100 dB-peSPL) with the grand-average individualized ABR simulations across the yNH group. Given that simulated ABR latencies were not impacted by CS, the applied latency shifts will not confound the CS prediction.

### Individual Synaptopathy Profile Predictions

In previous sections, cochlear model parameters of the subjects were determined using either AudTH- or DPTH-based methods and 13 personalized AEP-derived metrics were simulated for six CS profiles of each experiment participant. Here, we develop a classification approach, forward-backward classification, to predict the simulated CS profile that best matches recorded individual AEP metrics and determine the AEP metric that gives the most accurate segregation of simulated individualized CS profiles. This step was implemented separately for either of the cochlear individualization methods. After excluding eight subjects from the cohort (six without complete DPTHs and two with undetectable ABRs), we developed our individual SNHL-profiling method on data from 35 subjects (yNH: 12, oNH: 12 and oHI: 11).

Before classification, we first normalized the 13 AEP metrics (Table 1) derived from measured (M) and simulated six CS profiles per individual (S). The normalized S and M were calculated using Eq. 14 and 15.

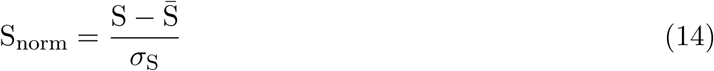

S is the matrix of simulated AEP metrics and contains 210 rows (35 subjects with six CS profiles) and 13 columns, the number of derived AEP metrics. deviation of S, respectively. 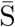 and *σ*_S_ refer to the mean and standard deviation of S, respectively.

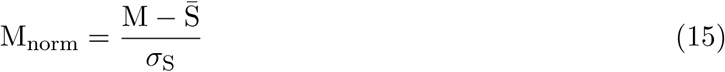

In Eq. 15, M refers to the matrix of measured AEP metrics with a dimension of 35*×*13. We created 8191 feature-sets using all possible combinations of 13 metrics 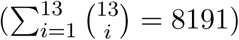. Metrics combination was performed separately for M_norm_ and S_norm_. The number of metrics in each feature-set varied between one and 13. From here on, **F** refers to the constructed 8191 feature-sets of AEP-derived metrics and F_i_ with *i* ∈ {1, …, 13} indicates a subset of **F** that has 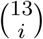 feature-sets and each feature-set contains a combination of i metrics. In the following paragraphs we explain the classification approach for an exemplary feature-set, f_e_, selected from **F**. The train and test datasets required for classification were constructed by choosing f_e_ of all participants from S_norm_ and M_norm_ and we called them S_train_ and M_test_, respectively. The proposed forward-backward classification method, comprised of two identical k-nearest-neighbor (kNN: k=1, Euclidean distance) classifiers. Classifier(1) in forward classification was trained by S_train_ in six classes with known class labels from the model simulations (L_S_), i.e. the six simulated CS profiles previously described in Fig. 3. Then, individual CS profiles were predicted by testing the trained classifier with the M_test_. Figure. 6a visualizes the different steps in forward classification. In this step, the evaluation of classification performance is unfeasible, since the actual CS degree of experiment participants are unknown. To address this issue, we interchanged the train-test datasets of the forward classification and implemented a second classification approach, called backward classification to assess the performance of the classifier(1) based on a second classifier (Fig. 6b). In this regard, we took the output of forward classification, i.e. the predicted CS degrees of experiment participants (L_M_ in Fig. 6), and corresponding measured AEP metrics (M_test_) to train the classifier(2) of Fig. 6b. Afterwards, S_train_, with known CS labels (L_S_) from the simulated individualized CS profiles, was used to test the trained classifier(2). The vector of predicted CS labels by classifier(2) (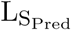) was compared to L_S_ and correspondig prediction accuracy was calculated as follows:

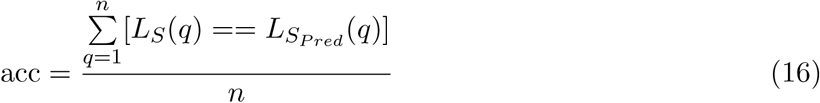

where *n* is equal to 210 (35 subjects with six CS profiles). Thus, the backward classification offers the possibility to calculate the accuracy of predicted CS profiles of study participants based on model simulations. We then repeated the forward-backward classification over all possible combinations of the derived metrics, i.e. 8191 feature-sets, and calculated the prediction accuracy of each feature-set according to Eq. 16. In this respect, the backward classification method gives the insight that to which degree classifier(1) was accurate in predicting CS degrees of experimental participants. Our classification approach makes use of combined simulated and recorded data to predict CS-profiles and can test the accuracy of these methods, even though a direct and actual validation of the CS histopathology still remains hidden due to experimental difficulties.

**Figure 6:**
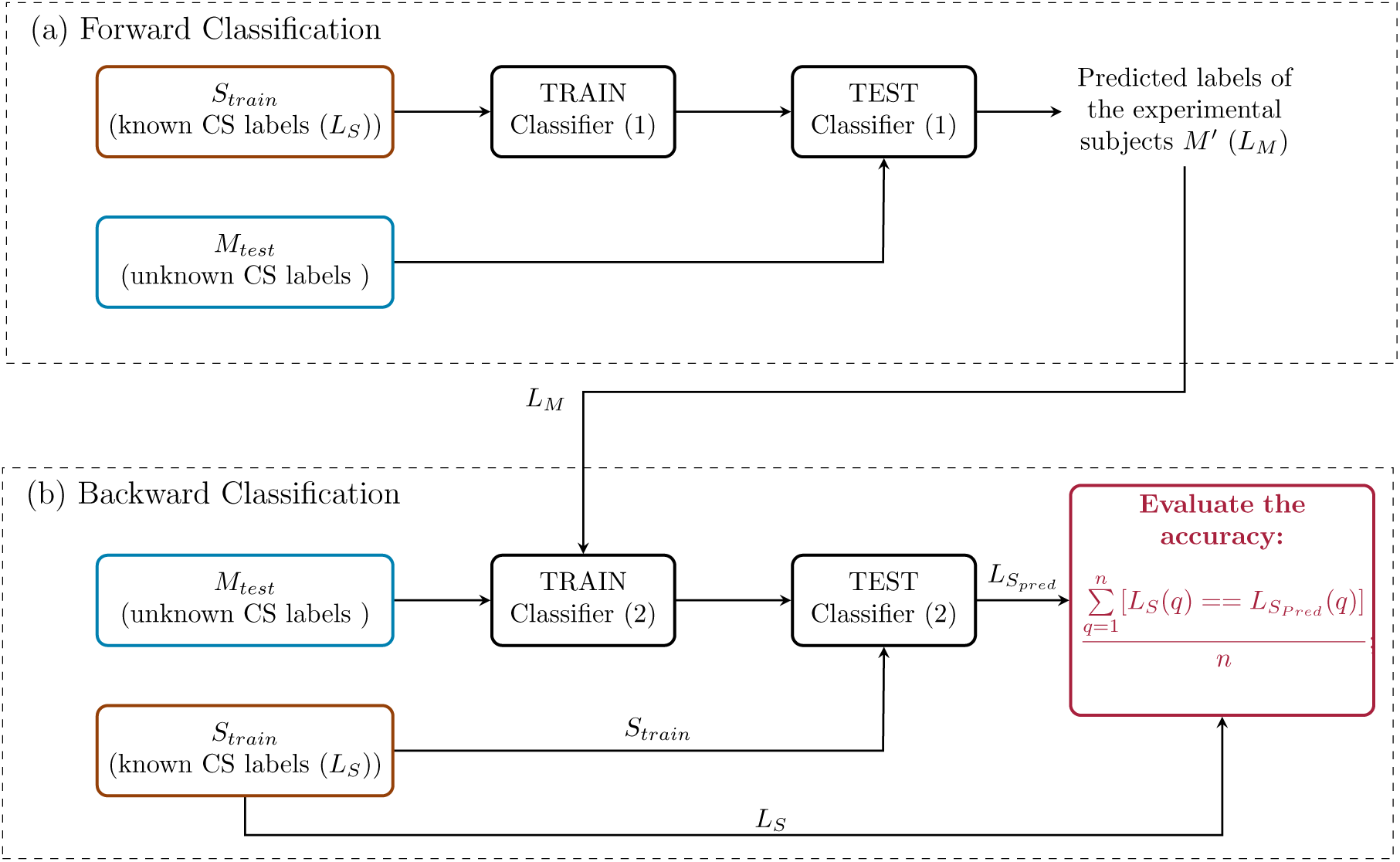
The forward-backward classification method. (a) Forward classification: Classifier(1) is trained with individualized simulated AEP-derived metrics (S_train_) for six CS profiles (L_S_) and tested with measured AEP-derived metrics (M_test_). The predicted labels (L_M_) for the study participants are entered to block (b). The backward-classification in (b) trains classifier(2) using measured AEP-derived metrics, i.e., M(test), and labels predicted by the forward classification i.e., L_M_. Classifier(2) is tested by S_train_ and corresponding labels (L_S_) are used to assess the classifier performance.

## Results

We applied forward-backward classification for each of the cochlear model individualization methods (AudTH and DPTH) and calculated the prediction accuracy of all feature-sets in **F**. For each cochlear profiling method, first, we determined the feature-set in each F_i_ (*i* ∈ {1, …, 13}), that had the highest classification accuracy. F_i_ consisted of feature-sets with *i* AEP-derived metrics. Then, the prediction variability was estimated using forward-backward classification by including the standard deviations of selected feature-sets. Lastly, we report individually predicted CS profiles belonging to those feature-sets.

### Combination of AEP-derived Metrics

To determine the best combination of metrics for CS profiling, the forward-backward classification was performed on the mean AEP-derived metrics of experiment participants and corresponding classification accuracy was reported as acc_mean_. Thus, we calculated acc_mean_ values of the predictions for 8191 feature-sets in **F** and determined the feature-set that yielded the highest acc_mean_ among all feature-sets in F_i_, with i combined metrics (*i* ∈ {1, …, 13}). Accordingly, 13 feature-sets were selected among 8191 in **F**. Table 2 and 3 list those feature-sets and corresponding acc_mean_ values for AudTH and DPTH-based methods, respectively. The RAM-EFR metric yielded the highest acc_mean_ values for both cochlear model individualization methods. The obtained 83.81% acc_mean_ of DPTH-based method, was higher than that of the AudTH-based method (68.57%), suggesting that methods which assess OHC damage more directly (i.e. DPTH vs. AudTH) yield a better classification accuracy in predicting simulated individualized CS profiles.

**Table 2:** Combination of metrics with the highest mean accuracy (acc_mean_) values in each F_i_, with *i* combined metrics. The standard deviation of obtained accuracies are shown in acc_sd_ column. The reported results are based on AudTH-based cochlear model individualization method.

| Involved Metrics | Involved Subjects | Best Combination of Metrics | acc (%) |  |
| --- | --- | --- | --- | --- |
| | | | $\text{acc}_{\text{mean}}$ | $\text{acc}_{\text{sd}}$ |
| 1 | 35 | RAM-EFR | 68.57 | 2.95 |
| 2 | 35 | RAM-EFR, w-$V_{\text{lat}100}$ | 64.76 | 1.73 |
| 3 | 35 | RAM-EFR, w-$I_{\text{lat}100}$, w-$I_{100}$ | 53.33 | 7.86 |
| 4 | 35 | RAM-EFR, w-$V_{\text{lat}100}$, w-$I_{\text{lat}100}$, w-$I_{100}$ | 51.90 | 9.28 |
| 5 | 35 | RAM-EFR, w-$V_{\text{lat}100}$, w-$I_{\text{lat}100}$, w-$I_{100}$, w-$V_{70}$ | 52.86 | 8.69 |
| 6 | 35 | RAM-EFR, w-$I_{\text{lat}100}$, w-$I_{100}$, w-$V_{70}$, w-$I_{70}$, w-V-growth | 51.43 | 6.97 |
| 7 | 35 | RAM-EFR, w-$V_{\text{lat}}\text{-growth}$, w-V-growth, w-$V_{\text{lat}100}$, w-$I_{70}$, w-$V_{70}$, w-$V_{100}$ | 45.24 | 6.79 |
| 8 | 35 | RAM-EFR, w-$V_{\text{lat}}\text{-growth}$, w-V-growth, w-$V_{\text{lat}100}$, w-$V_{\text{lat}70}$, w-$I_{70}$, w-$V_{70}$, w-$V_{100}$ | 45.24 | 6.59 |
| 9 | 35 | RAM-EFR, w-V-growth, w-I-growth, w-$I_{\text{lat}100}$, w-$V_{\text{lat}100}$, w-$I_{70}$, w-$V_{70}$, w-$I_{100}$, w-$V_{100}$ | 36.19 | 7.11 |
| 10 | 35 | RAM-EFR, w-V-growth, w-I-growth, w-$V_{\text{lat}}\text{-growth}$, w-$V_{\text{lat}100}$, w-$V_{\text{lat}70}$, w-$I_{70}$, w-$V_{70}$, w-$I_{100}$, w-$V_{100}$ | 32.86 | 6.67 |
| 11 | 35 | RAM-EFR, w-V-growth, w-I-growth, w-$V_{\text{lat}}\text{-growth}$, w-$I_{\text{lat}}\text{-growth}$, w-$V_{\text{lat}100}$, w-$V_{\text{lat}70}$, w-$I_{70}$, w-$V_{70}$, w-$I_{100}$, w-$V_{100}$ | 27.62 | 6.49 |
| 12 | 35 | RAM-EFR, w-V-growth, w-I-growth, w-$V_{\text{lat}}\text{-growth}$, w-$I_{\text{lat}}\text{-growth}$, w-$V_{\text{lat}100}$, w-$V_{\text{lat}70}$, w-$I_{70}$, w-$V_{70}$, w-$I_{100}$, w-$V_{100}$, w-$I_{\text{lat}70}$ | 18.10 | 6.65 |
| 13 | 35 | RAM-EFR, w-V-growth, w-I-growth, w-$V_{\text{lat}}\text{-growth}$, w-$I_{\text{lat}}\text{-growth}$, w-$V_{\text{lat}100}$, w-$V_{\text{lat}70}$, w-$I_{70}$, w-$V_{70}$, w-$I_{100}$, w-$V_{100}$, w-$I_{\text{lat}70}$, w-$I_{\text{lat}100}$ | 17.14 | 6.75 |

**Table 3:**
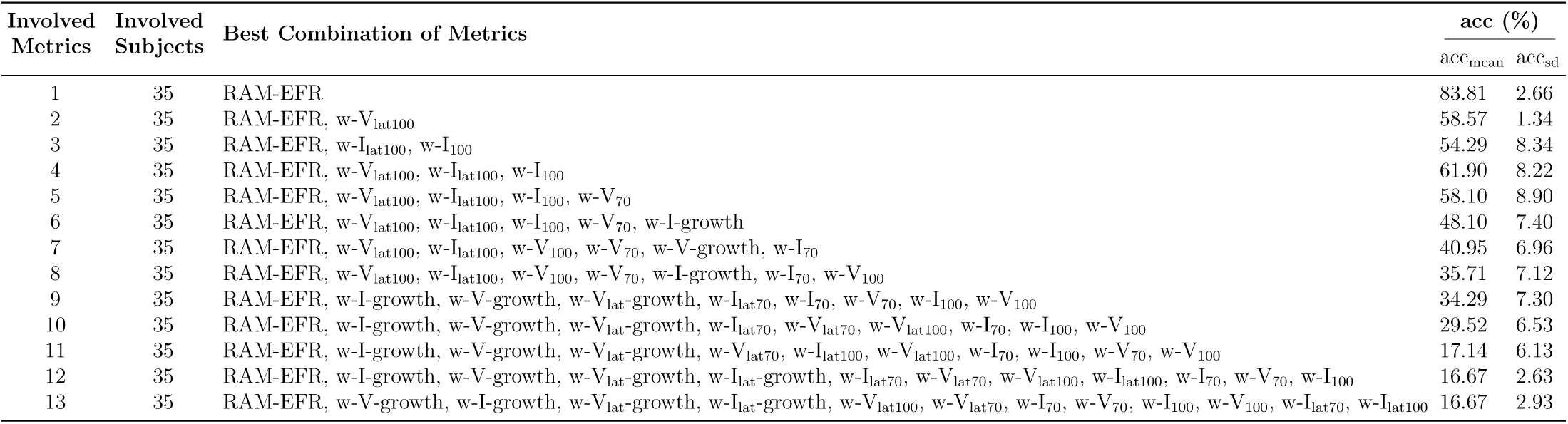
Combination of metrics with the highest mean accuracy (acc_mean_) values in each F_i_, with *i* combined metrics. The standard deviation of obtained accuracies are shown in acc_sd_ column. The reported results are based on DPTH-based cochlear model individualization method.

### Prediction Variability

The impact of subject-specific factors and measurement noise reflect on inter- and within-subject variability of the AEP recordings and can have an impact on the accuracy of the classification method. To measure this effect, the forward-backward classification was repeated, this time by extracting metrics from the bootstrapped average trials, rather than from the mean of trials. This resulted in distributions for each specific metric and each subject, with standard deviations as given by the last column of Table 1. Then, 100 samples were randomly drawn from the distribution of each metric. Thus, for every feature-set in Table 2 and 3, the corresponding metrics samples were combined to yield 100 variations of each feature-set. Afterwards, the CS profile prediction was repeated 100 times with each feature-set for each subject, and prediction accuracy was assessed in every repetition. Lastly, the standard deviation of the calculated accuracies (acc_SD_) was determined over the 100 repetitions of each feature-set and listed in the last column of Table 2 and 3.

For the best predictor metric (RAM-EFR), acc_SD_ values of 2.95% and 2.66% were obtained for the AudTH- and DPTH-based methods, respectively. The lowest acc_SD_ was obtained when combining the RAM-EFR with the w-V_lat100_ metric in both cochlear model individualization methods (AudTH: 1.73% and DPTH: 1.34%). However, the respective acc_mean_ values were considerably lower than those of the RAM-EFR by itself, particularly in DPTH-based method. To assess the performance of the RAM-EFR based CS profile prediction in sub-groups, we show confusion tables in Fig. 7 for AudTH- and DPTH-based cochlear model individualization methods. The diagonals of each table reflect how often the classifier assigned a CS profile (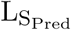: predicted class) that matched with that of in simulated individualized CS profiles (L_S_: true class). Off-diagonal values show the number of instances that 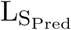 and 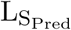 were not identical. Detailed prediction accuracy values of each sub-group are summarized in the tables in Fig. 7. The highest and lowest prediction accuracy values relate to the yNH and oHI group, respectively for both AudTH- and DPTH-based methods. Comparing the cochlear model individualization methods, it is seen that the DPTH-based approach outperforms the AudTH-based method on both group- and sub-group levels.

**Figure 7:**
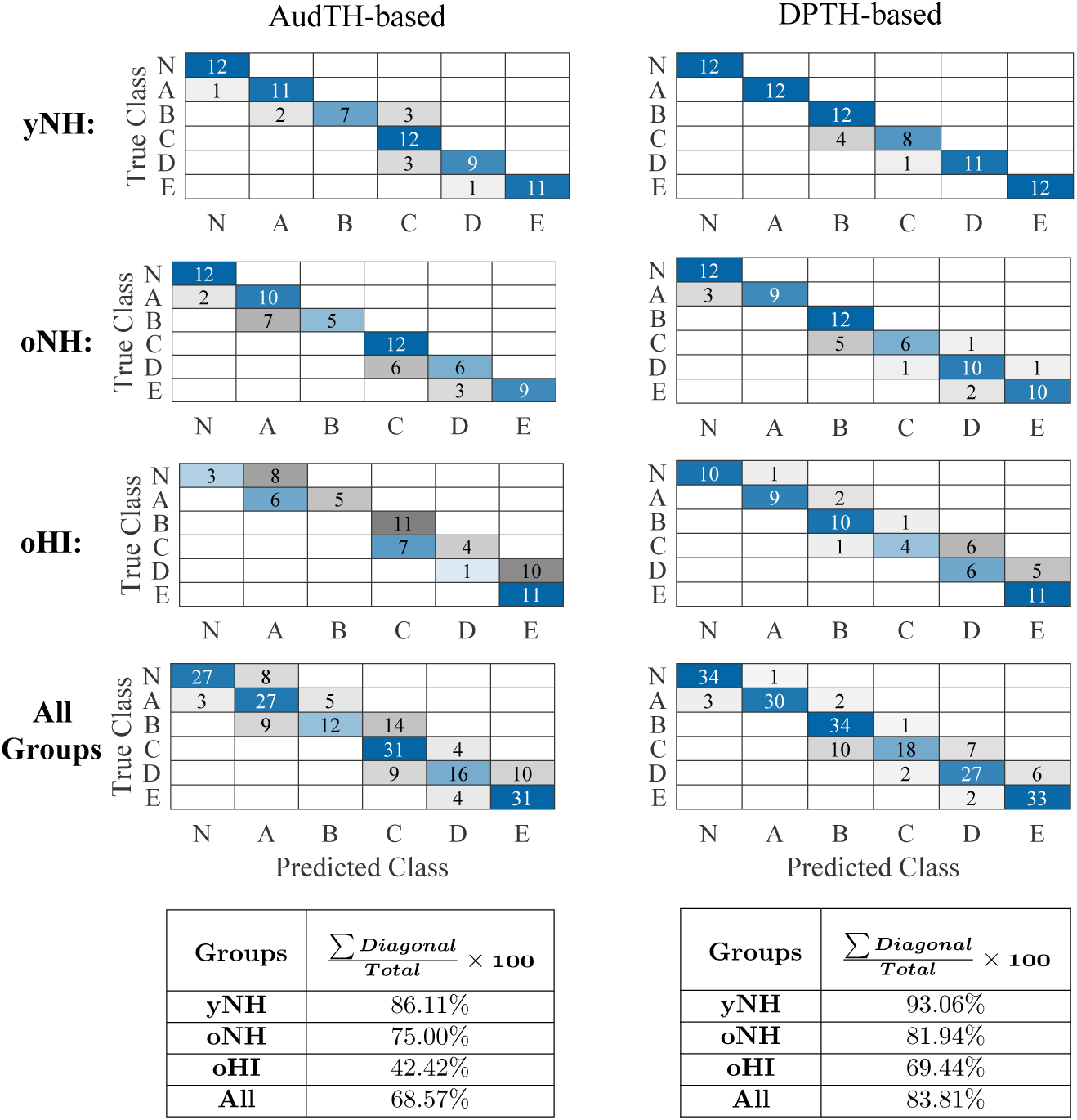
Confusion tables at subgroup and group-levels for both AudTH and DPTH-based cochlear model individualization methods. The tables summarize the accuracy of classifier(2) in Fig. 6b for subgroups, as well as all groups together.

### Cochlear synaptopathy profile prediction based on individualized classifiers

Table 4 lists the predicted individual CS profiles from the RAM-EFR metric (best prediction accuracy) for both AudTH- and DPTH-based cochlear individualization methods. The reported profiles are the output of the forward classification step, i.e. L_M_ shown in Fig. 6. Considering the AudTH and DPTH columns of Table 4, lower degrees of AN-damage were predicted for the yNH group than for the oNH and oHI groups. Additionally, the range of predicted CS profiles in the yNH group shows that yNH listeners might also suffer from different degrees of CS. The oHI group, which was assumed to suffer from mixed OHC-damage and CS pathologies, were predicted to have the highest degree of CS among the cohort.

**Table 4:** Predicted individuals CS profiles obtained from AudTH and DPTH based cochlear individualization methods, based on RAM-EFR metric. Columns AudTH_ind_ and DPTH_ind_, list the predicted CS profiles by desinging individualized classifiers based on RAM-EFR metric.

| Group | No. | AudTH | AudTH<sub>ind</sub> | DPTH | DPTH<sub>ind</sub> |
| --- | --- | --- | --- | --- | --- |
| yNH | 1 | C | B | B | B |
|  | 2 | A | A | A | B |
|  | 5 | N | N | N | N |
|  | 7 | N | N | N | N |
|  | 8 | N | N | N | N |
|  | 9 | N | N | N | N |
|  | 10 | N | N | N | A |
|  | 11 | A | B | B | B |
|  | 12 | N | N | N | A |
|  | 13 | A | A | A | A |
|  | 14 | N | N | N | N |
|  | 15 | N | N | N | N |
| oNH | 1 | D | D | C | D |
|  | 3 | E | E | E | E |
|  | 4 | D | E | D | D |
|  | 6 | D | D | D | D |
|  | 7 | C | D | D | D |
|  | 8 | E | E | E | E |
|  | 9 | N | A | N | A |
|  | 10 | B | B | B | B |
|  | 11 | C | D | D | D |
|  | 12 | N | N | N | N |
|  | 13 | E | E | E | E |
|  | 14 | C | D | C | C |
| oHI | 1 | E | E | E | E |
|  | 2 | E | D | E | D |
|  | 3 | E | E | E | E |
|  | 4 | E | E | E | E |
|  | 5 | E | D | E | E |
|  | 7 | E | D | E | E |
|  | 8 | E | E | E | E |
|  | 9 | E | E | E | E |
|  | 10 | E | E | E | E |
|  | 11 | E | E | E | E |
|  | 12 | E | E | E | E |

Thus far, the reported individualized CS profiles for RAM-EFR were predicted by training a single classifier with simulated individualized CS profiles of the whole experimental cohort. This has drawbacks for individual profiling in a clinical context, because it would be ideal if the profiling could be performed using only recordings from the tested individual. Hence, to establish more accurate predictions of the individual CS degrees, we took one step further and designed individualized classifiers, which were trained and tested with the RAM-EFR metric of the same listener. If RAM_s_ stands for the six simulated CS profiles of a nominal subject and RAM_m_ for the measured RAM-EFR metric, we first normalized RAM_s_ and RAM_m_ values by the 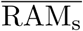 and 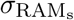 (mean and standard deviation of RAM_s_). Then we trained and tested the classifier, with the same characteristics as classifier(1) and (2), using normalized RAM_s_ and RAM_m_ values, respectively. This procedure was repeated for all listeners in the cohort and for both AudTH and DPTH-based cochlear model pole-setting methods. The predicted individualized CS profiles were listed in Table 4 (columns: AudTH_ind_ and DPTH_ind_). Considering either of the AudTH- or DPTH-based methods, designing individualized classifiers revealed only minor differences in the predicted CS profiles of individual listeners compared to those predicted by a single classifier trained with simulated individualized RAM-EFRs. However, the CS profiles reported in AudTH_ind_ and DPTH_ind_ columns might be more reliable than the group-based predictions, since the former were predicted by individualized classifiers that were trained on the basis of personalized cochlear simulations.

To provide a demonstration of the implemented method, and to show to which extent the model simulations imitate the experimental measurements, we compare simulated and measured AEPs of a yNH subject in Fig. 8. Panel (a) depicts simulated RAM-EFR spectra for the different considered CS profiles. Based on the experimental RAM-EFR (panel (d)) and forward classification, we predicted that this subject had a “N” CS profile, i.e., no AN-damage. The CGL parameters of the individualized model were adjusted based on DPTHs of the same yNH listener. Panels (b) and (c), depict the simulated personalized ABR waveforms for the predicted “N” CS degree. Experimental ABR waveforms to 70 and 100 dB-peSPL clicks are shown in panels (e) and (f), respectively. Details regarding the value of extracted metrics from the measurements and simulations are provided in a table at the bottom of the Fig. 8. Even though our classifier did not consider ABR metrics, the applied personalized OHC and AN profiles predicted w-I_lat100_, w-I_70_, w-V_70_ and w-I_100_ markers that fell the standard deviation of the corresponding recorded values. The remaining simulated ABR metrics, i.e., w-I_lat70_, w-V_lat100_ and w-V_100_, only minimally deviated from the range of respective measurements, showing that our method accurately predicts AEP features to stimuli which were not included in the classifier.

**Figure 8:**
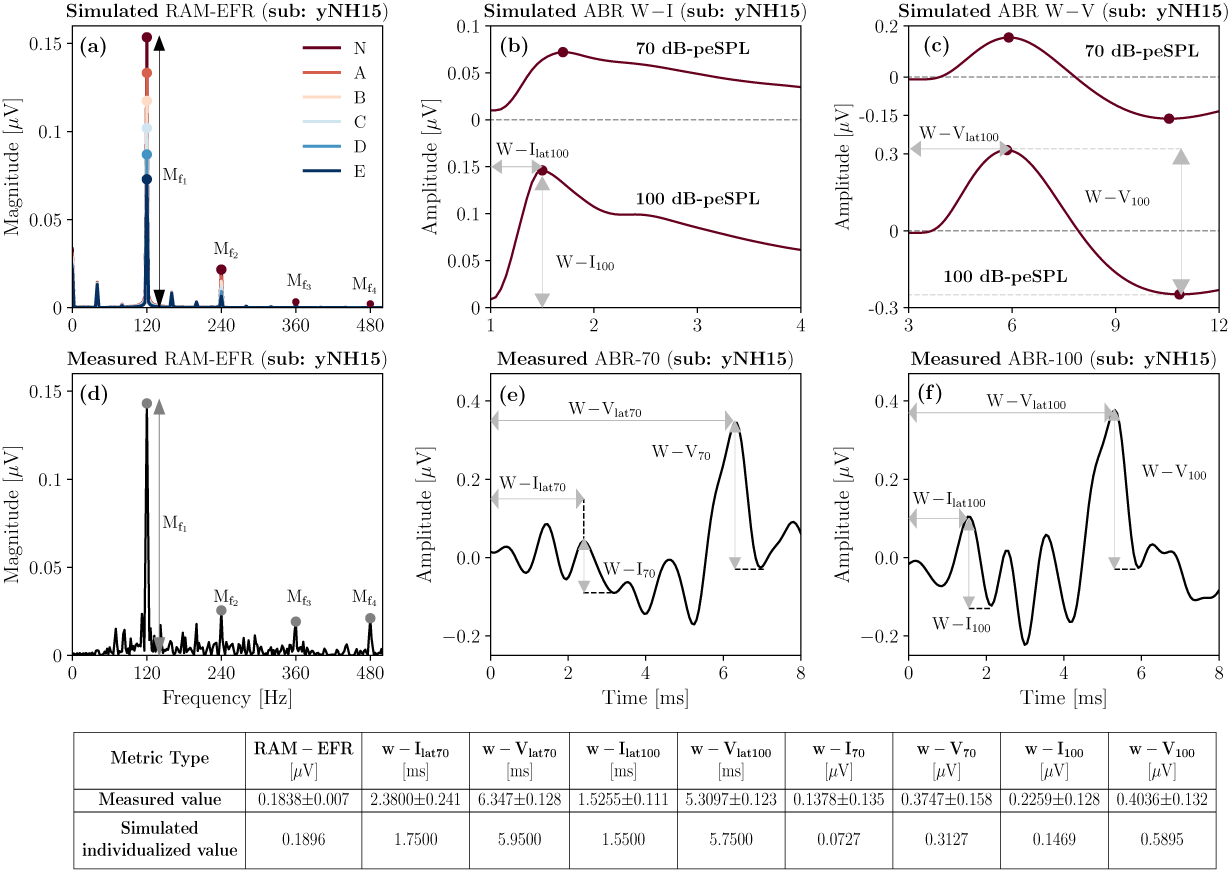
A comparison between simulated and measured AEPs for a yNH subject (yNH15). This subject was predicted to have a normal (N) CS profile, i.e., without CS. (a) Simulated RAM-EFR spectra for six CS profiles. The sum of the drawn arrows yields the RAM-EFR magnitude metric. (b) Simulated ABR wave-I to 70 and 100 dB-peSPL clicks. Waveforms were shifted by 1ms to match the experimental data. (c) Simulated ABR wave-V to 70 and 100 dB-peSPL clicks. Waveforms were shifted by 3ms to match the experimental data. The specified arrows in (b) and (c) indicate the extracted metrics. (d) Measured RAM-EFR of the same listener (yNH15). Shown arrows, indicate the peak-to-noisefloor values. Akin to (a), the measured RAM-EFR metric was calculated by summing the of arrow amplitudes. (e) Measured ABR waveform to 70 dB-peSPL clicks. (f) Measured ABR waveform to 100 dB-peSPL clicks. Arrows in (e) and (f), determine the extracted metrics. The shown simulated waveforms were predicted based on the DPTH-based cochlear individualization method. The table shows the exact value of EFR and ABR metrics derived from recordings and predicted CS-profile, “N”, of the same listener.

### Method Validation

To validate the proposed method and its generalizability to other cohorts and other measure- ment equipment, we applied the developed classifier in backward classification step to RAM-EFRs recorded in a second experiment. Figure 9 schematizes the implementation of the validation method. Considering the different experimental setup and recording location of the second experiment, the measured RAM-EFRs of both experiments were scaled between zero and one, prior to classification. Given that only yNH listeners participated in the second experiment, we employed the smallest RAM-EFR magnitude recorded from oHI listeners (as part of another study) recorded with the same setup as the second experiment to scale the RAM-EFRs. The scaled RAM-EFRs of the first experiment were used to train the classifier(1) in Fig. 6 and afterwards, the trained classifier was tested with the scaled RAM-EFRs of the second experiment. The predicted CS profiles are illustrated as a bar-plot in Fig. 9. 84.21% of the 19 yNH participants of the second experiment were classified as *N*, i.e. without CS, and the rest were predicted to have mild CS. These predictions show that a classifier designed on our cohort can be applied to other cohorts to predict individual CS degrees based on the RAM-EFR. In line with expectations, the classifier predicted that most yNH subjects were synaptopathy free.

**Figure 9:**
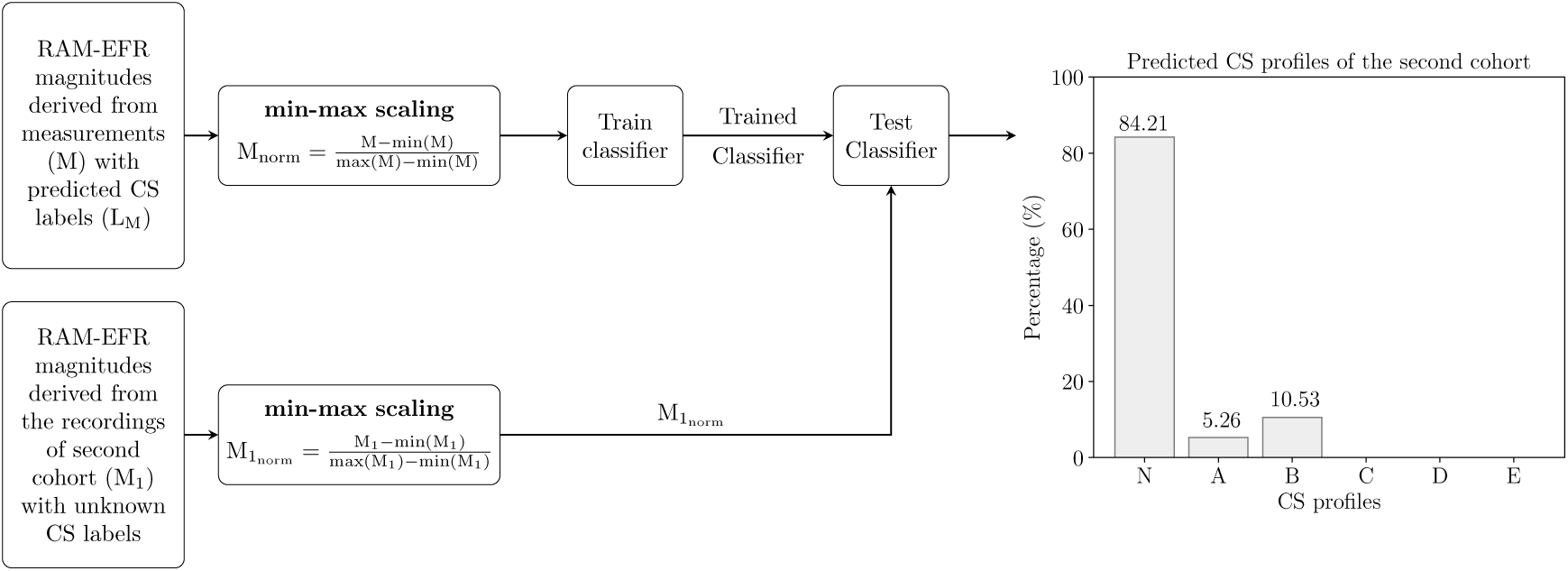
Implementation of the validation method. Measured RAM-EFRs (M) with predicted labels in Fig. 6 (L_M_) are scaled between zero and one to train a kNN classifier. The trained classifier is tested with scaled RAM-EFRs recorded from the second cohort comprised of yNH listeners. The bar-plot shows the predicted CS profiles for the second cohort listeners. The CS profiles labels in the bar-plot are similar to those defined in Fig. 3.

## Discussion

By combining experimental ABR and EFR measurements with a modelling approach, we were able to develop a classifier that can assign one out of six CS profiles to listeners with mixed SNHL pathologies. The classifier considered 8191 feature-sets, of which our forward-backward classification method identified that the RAM-EFR metric yielded the best performance in both AudTH- and DPTH-based cochlear individualization methods. We tested both a group and individually based method and showed that our method can generalize to other cohorts and measurement setups. Taken together, we have high hopes that this method can find its way to clinical hearing diagnostics, since a single AEP metric is required to yield a CS-profile prediction, given the audiogram or at least four DPTHs.

### Implications for RAM-EFR-based synaptopathy profiling prediction

On the one hand, predicting the CS degree from AEP metrics is controversial in listeners with coexisting OHC deficits and on the other, validation of the predicted CS profiles with temporal bone histology is impossible in humans. Without these means, models of the human auditory periphery an AEP generators can provide a tool to bridge this experimental gap. The similarity between predicted AEP degradations for a known CS profile and experimental AEP degradations can be used to predict the CS profile of individuals. In a previous study, we tested the potential of the derived-band EFR as a CS predictor in NH listeners using a fuzzy c-means clustering method, and validated our CS predictions using an another AEP-derived metric (wave-V amplitude growth slope) recorded from the same listener. We evaluated the method based on the percent of subjects that were predicted and validated to have the same CS profile, i.e. 61% (Keshishzadeh and Verhulst, 2019). However, the performance of this method is easily impacted by the characteristics of the adopted predictor and validation metrics, e.g. different generator sources, degree of sensitivity to subtypes of SNHL and tonotopic susceptibility.

The interdisciplinary approach we took in this study tackled this validation issue by proposing a forward-backward classification approach and applying the trained classifier to AEPs from a new cohort to test its generalizability. Moreover, we were able to determine the most accurate AEP-derived metric for CS degree prediction, given a range of 13 possible AEP-derived metrics. Among the considered AEP-based metrics and combinations thereof, we found that the RAM-EFR magnitude showed the best performance in segregating simulated individual CS profiles. At the same time, RAM-EFR metric was involved in all feature-sets that yielded the highest acc_mean_ among feature-sets that had equal number of combined metrics (Table 2 and 3). This finding is consistent with the outcome of Parthasarathy and Kujawa (2018) and Vasilkov et al. (2020), showing that EFRs to SAM or RAM are sensitive to CS. Moreover, the combined modelling and experimental study of Vasilkov et al. (2020) showed that the adopted RAM-EFR marker (RAM with a 25% duty-cycle), is minimally impacted by OHC damage. The sharp envelope combined with the long silence intervals between stimulus peaks generates more synchronized AN fiber responses compared to conventional SAM stimulus to yield a stronger EFR with extended dynamic range across subjects. Lastly, the RAM-EFR is a more sensitive marker of CS than ABR (Parthasarathy and Kujawa, 2018). Taken together, our results indicate that the RAM-EFR magnitude is an appropriate AEP-based metric to predict individual CS degree of listeners in the presence of OHC-loss.

### The effect of cochlear model individualization method on predicting cochlear synaptopathy profiles

In this study, we determined the CGL model parameters using either measured audiometric or DPOAE thresholds, and assessed the classifier performance of each method in the backward classification step. Comparing the resulting acc_mean_ values for each cochlear individualization method can inform which of the two methods yielded the most accurate AEP simulations for a given CS profile. The acc_mean_ values of RAM-EFR metric showed that setting cochlear filter pole functions on the basis of measured DPTHs outperforms the AudTH method for all experimental groups (Fig. 7, Tables 2 and 3). This outcome is consistent with literature studies showing that OAEs are a more sensitive measure of noise-induced cochlear dysfunction in humans (Engdahl et al., 1996; Konopka et al., 2005; Seixas et al., 2005; Marshall et al., 2009). Moreover, OAEs are not influenced by inner-hair-cell/AN damage (Trautwein, 2002), whereas behaviourally measured audiometric thresholds, particularly extended high-frequency thresholds, could be affected by extreme neural degeneration (Lobarinas et al., 2013; Liberman et al., 2016; Bramhall et al., 2019). Consequently, given the varied susceptibility of AudTHs and DPOAEs to different aspects of SNHL, it was expected that we would obtain non-identical predictions of CS profiles for a nominal subject (Table 4). Comparing the AudTH and DPTH columns in Table 4, we found a mismatch between individually predicted CS profiles for 14.28% of subjects (yNH: two, oNH: three). The mismatch degree increased to 20% (yNH: three, oNH: two and oHI: two) when the individual CS profiles were predicted using personalized classifiers (AudTH_ind_ and DPTH_ind_ columns).

It is noteworthy that the DPTH-based cochlear individualization was implemented using DPTHs from only four frequencies (0.8 to 4 kHz), whereas the AudTH-based method considered audiometric thresholds measured at 12 frequencies (0.125 to 10 kHz). This difference may have resulted in less accurate CGL model parameters for the DPTH-based method, despite a better performance of forward-backward classification. In future implementations of this method, we intend to incorporate more frequencies in the DPTH measurements, especially at higher frequency regions. Employing DP-grams instead of DPTHs is another option, as these require a shorter measurement time. In both cases, we suggest to include lower stimulus levels as well, given that noise-induced OHC deficits can be identified earlier at lower stimulus levels (Bramhall et al., 2019).

## Method Limitations

The proposed method for AEP-based CS-profiling, relies on the interactive use of recordings and model simulations. Hence, shortcomings in either aspect could have caused performance limitations of the method. The following sections summarize a number of these limitations:

### Experimental Limitations

(1) ABRs in humans are recorded using vertex electrodes placed on the scalp, which yields smaller and more variable wave-I amplitudes than when they are recorded in animals using subdermal electrodes. The measured ABR w-I_70_ amplitude in our measurement produced a mean standard deviation of 0.198*µ*V across the cohort, which is fairly large with respect to the mean amplitude of 0.146*µ*V (yNH: 0.1964±0.1436*µV*, oNH: 0.1304±0.203*µV*, oHI: 0.1071±0.243*µV*). Compared to w-I_70_, w-I_100_ amplitudes showed less variability, i.e., 0.2503±0.2056*µV*. Variability of the w-I_100_ was considerably lower only for yNH group (0.350±0.143*µV*). Per subgroup, variability increased for older groups (oNH: 0.205±0.247*µV*, oHI: 0.180±0.235*µV*). Given these variabilities, adding the w-I_100_ metric to the second feature-set (RAM-EFR, w-V_lat100_), suddenly increased the acc_SD_ (Tables 2 and 3). (2) Although adopting relative ABR metrics, such as growth functions might factor out individual differences, the standard deviation of the derived relative metric is influenced by the propagated error of the absolute metric. (3) ABRs to clicks presented at 100 dB-peSPL should yield higher wave-I and V amplitudes, than when the stimulus was presented at 80 dB-peSPL. Nevertheless, the opposite was observed in a few subjects.

### Model Limitations

(1) The adopted computational model of the auditory periphery allows for OHC deficit simulation on a CF-dependent basis, but not for CGLs above 35 dB, since the maximum possible BM filter gain is 35 dB in the model (Verhulst et al., 2018). This constraint led to elevated absolute prediction errors for high-frequency audiometric thresholds in the oHI (above 4 kHz) and oNH (above 8 kHz) groups (Fig. 4e). The increased absolute errors were mainly observed for the audiometric threshold predictions, since DPTHs were only measured for frequencies up to 4 kHz. Thus, the individualized hard-coded OHC-loss component for the oHI group might lead to similar and less accurate CS profile prediction for oHI participants with audiometric losses greater than 35 dB-HL. (2) In the adopted method, we hard-coded the CGL using the individual hearing thresholds and related the remaining AEP alterations to CS. An alternative way would be to run the model iteratively and simultaneously optimize both CGL and CS profile parameters on the basis of the experimental data to obtain the best OHC-loss and CS profiles. However, we did not further explore this route due to the high computational cost of running the adopted TL cochlear model in an iterative optimization procedure.

## Conclusion

In this study, we proposed an integrated modelling and experimental approach to build personalized auditory models and predict the AN-damage profile of listeners with mixed SNHL profiles. To develop individualized cochlear models, we implemented two different methods on the basis of measured AudTHs and DPTHs. Next, we developed a classification-based approach to predict individual CS profiles and determined which AEP metric (or combinations thereof) yielded the highest prediction accuracy. Afterwards, we evaluated the implemented CGL and CS-profile individualization methods on the development dataset, as well as on a new cohort. Our study suggests that a DPTH-based cochlear model individualization approach combined with a RAM-EFR recording predicts individual CS profiles most accurately among the 8191 possible combinations of 13 AEP markers. Additionally, we tested the applicability of the proposed method by applying the trained classifier to the recorded RAM-EFRs of a new cohort of yNH listeners. The classifier predicted that these listeners mostly had mild forms of CS, which supports that our method is generalizable to other recording setups and cohorts. Training the classifier again on larger cohorts may further increase the generalizability of the method. We hope that this method, or variations thereof, can be used in a clinical diagnostic context, as the number of needed AEP recordings to yield an individual CS-profile is small (i.e. 10-15 minutes). Individualized models of SNHL are an important step for the development of hearing aid algorithms that compensate for both the OHC- and AN-damage aspects of SNHL.

## Acknowledgement

This work was supported by European Research Council (ERC) under the Horizon 2020 Research and Innovation Programme, grant agreement No. 678120 RobSpear (SK, SV) and DFG Cluster of Excellence EXC 1077 1 Hearing4all (MG, SV).

## Notes

### Competing Interest Statement

The authors have declared no competing interest.

